# Phytochrome B regulates reactive oxygen signaling during abiotic and biotic stress in plants

**DOI:** 10.1101/2021.11.29.470478

**Authors:** Yosef Fichman, Haiyan Xiong, Soham Sengupta, Rajeev K. Azad, Julian M. Hibberd, Emmanuel Liscum, Ron Mittler

## Abstract

Plants are essential for life on Earth converting light into chemical energy in the form of sugars. To adjust for changes in light intensity and quality, and to become as efficient as possible in harnessing light, plants utilize multiple light receptors, signaling, and acclimation mechanisms. In addition to altering plant metabolism, development and growth, light cues sensed by some photoreceptors, such as phytochromes, impact on many plant responses to biotic and abiotic stresses. Central for plant responses to different stresses are reactive oxygen species (ROS) that function as key signaling molecules. Recent studies demonstrated that respiratory burst oxidase homolog (RBOH) proteins that reside at the plasma membrane and produce ROS at the apoplast play a key role in plant responses to different biotic and abiotic stresses. Here we reveal that phytochrome B (phyB) and RBOHs function as part of a key regulatory module that controls ROS production, transcript expression, and plant acclimation to excess light stress. We further show that phyB can regulate ROS production during stress even if it is restricted to the cytosol, and that phyB, RBOHD and RBOHF co-regulate thousands of transcripts in response to light stress. Surprisingly, we found that phyB is also required for ROS accumulation in response to heat, wounding, cold, and bacterial infection. Taken together, our findings reveal that phyB plays a canonical role in plant responses to biotic and abiotic stresses, regulating ROS production, and that phyB and RBOHs function in the same pathway.

**Significant Statement:** Abiotic and biotic stresses cause extensive losses to agricultural production and threaten global food security. Augmenting plant resilience to stressful conditions requires understanding of how plants sense stress. Here we report that the sensing of different abiotic and biotic stresses that result in the production of the key stress-response signaling molecules, reactive oxygen species, requires the plant photoreceptor protein phytochrome B. We further show that in contrast to its many nuclear functions, phytochrome B regulates reactive oxygen production by plasma membrane-localized respiratory burst oxidase homologs while localized to the cytosol. Our findings reveal the existence of a rapid stress response regulatory mechanism requiring phytochrome B and reactive oxygen species, essential for plant acclimation to stress.

## Introduction

Light is indispensable for plants serving as an energy source, developmental signal, and regulator of many different responses to environmental cues and stresses. However, light can come in different intensities and qualities that can rapidly fluctuate. To hone their responses to light, and to become as efficient as possible in harnessing it, plants utilize several different light receptors and signal transduction pathways that control multiple biochemical, physiological, metabolic, and molecular mechanisms and allow them to successfully adapt to changes in light conditions (1–7). While light energy is harvested by chloroplasts, that can independently adjust to changes in light conditions using different biochemical mechanisms (8–12), changes in light intensity and quality are sensed by light receptors, such as phytochromes (phys), cryptochromes (crys), UV-B resistance 8 (UVR8), ZEITLUPE (ZTL) and other proteins that regulate transcriptomics responses of plants (6, 13). In addition to regulating photosynthesis and linking light cues with different developmental and environmental responses, some of the biochemical and regulatory pathways described above evolved to prevent overloading of the photosynthetic apparatus during conditions of excess light stress, that could result in the unregulated production of potentially damaging reactive oxygen species (ROS; 14–17).

Multiple studies demonstrated that under conditions of excess light stress singlet oxygen (^1^O_2_) is produced by photosystem (PS) II and superoxide (O_2_·^-^) is produced by PSI in chloroplasts (14, 15, 18). The O_2_·^-^ produced by PSI dismutates to H_2_O_2_ (spontaneously or via superoxide dismutases; SODs), and H_2_O_2_ that is not scavenged inside the chloroplast is thought to diffuse into the cytosol through aquaporins (14, 15, 18, 19). In addition, H_2_O_2_ is produced in peroxisomes during photorespiration, and O_2_·^-^ and H_2_O_2_ can form in mitochondria as part of a retrograde signaling pathway triggered by excess light stress (20–23). The production of ROS by these different compartments during excess light stress is proposed to trigger different acclimation and retrograde mechanisms, as well as to activate different ROS scavenging mechanisms (14, 24).

As described above, the majority of ROS accumulating in plant cells during excess light stress were traditionally thought to originate in chloroplasts and mitochondria, and in C3 plants, in peroxisomes as well (14–17, 20–22, 24). However, new findings in Arabidopsis (*Arabidopsis thaliana*) and rice (*Oryza sativa*) revealed that during excess light stress, the majority of ROS that accumulate in plant cells are produced at the apoplast by RESPIRATORY BUST OXIDASE HOMOLOGs (RBOHs; 25, 26). This more recent work suggests that during excess light stress O_2_·^-^ and H_2_O_2_ are produced in the apoplast, rather than the chloroplast, mitochondria and/or peroxisomes, and that H_2_O_2_ diffuses from the apoplast into the cytosol via aquaporins and alters the redox state of different proteins triggering signal transduction pathways. Alternatively, ROS produced in the apoplast could trigger signal transduction pathways in the cytosol via plasma membrane (PM)-localized receptors such as CYSTEINE-RICH RECEPTOR-LIKE KINASEs (CRKs) and HYDROGEN-PEROXIDE-INDUCED Ca^2+^ INCREASE 1 (HPCA1; 27, 28). The highly regulated function of RBOHs, which generate O_2_·^-^ (that subsequently dismutates to H_2_O_2_) at the apoplast in response to many different developmental-, abiotic-, mechanical-, and pathogen-derived signals (29, 30), is therefore utilized for ROS production during excess light stress. Moreover, this process was shown to occur during excess light stress even in etiolated plants that contain undeveloped chloroplasts that do not conduct photosynthesis (25). These tantalizing results highlight a new question however of how is light sensed during excess light stress to trigger this apoplastic ROS production response?

A candidate receptor for light sensing that could lead to ROS production by RBOHs during excess light stress is phyB. In Arabidopsis and tomato (*Solanum lycopersicum*) phyB was found to be required for systemic stomatal responses, systemic ROS accumulation, and systemic photosynthetic regulation, in response to a local treatment of high light stress (26, 31). Here we report that phyB and RBOHs function as part of a key regulatory module that controls ROS production, transcript expression, and plant acclimation in response to excess light stress. We further show that phyB is required for ROS production during excess light stress even if it is restricted to the cytosol, and that phyB, RBOHD and RBOHF co-regulate 1000s of transcripts in response to excess light stress. Remarkably, we found that phyB is also required for ROS accumulation in response to heat, wounding, cold, and bacterial infection. Taken together, our findings suggest that phyB plays a pivotal role in plant responses to biotic and abiotic stresses, regulating ROS production, and that phyB and RBOHs likely function in the same regulatory pathway.

## Results

### PhyB, RBOHD and RBOHF regulate ROS accumulation, physiological responses, and plant acclimation to excess light stress

To study the role of phyB in ROS production during excess light stress, we subjected wild type (WT) plants and *phyB*, *rbohD*, *rbohF,* and *rbohD rbohF* mutants to an excess light stress treatment (740 μmole photons s^−1^ m^−2^) for 10 min and measured ROS accumulation in whole plants grown in soil using our newly developed live ROS imaging method (32–34). This method uses the cell-permeant 2’,7’-dichlorodihydrofluorescein diacetate (H_2_DCFDA) that detects a wide range of different ROS including H_2_O_2_ (33). In addition to white light, we also subjected plants to red light (120 μmole photons s^−1^ m^−2^) using a light intensity that is proportional to the fraction of red light included within our white light treatment. As shown in Fig. 1A, treatment with excess white or red light resulted in ROS accumulation in WT plants, but not in *phyB*, *rbohD*, *rbohF*, or *rbohD rbohF* mutants. To study the impact of excess light stress on ROS production in plants that do not contain photosynthetically active chloroplasts, we used etiolated seedlings. As shown in Fig. 1B, treatment of etiolated WT seedlings with white or red light resulted in ROS accumulation, while treatment of etiolated *phyB*, *rbohD*, *rbohF,* or *rbohD rbohF* seedlings did not. Interestingly, the amount of ROS produced in mature green plants treated with white light was higher than that of mature green plants treated with red light, while this difference was not observed in etiolated seedlings (Figs. 1A, 1B). This finding suggests that some ROS could be produced by photosynthetically active chloroplasts of mature green plants in response to white light. However, even in green plants, this amount of ROS production was completely abolished in the *rbohD rbohF* double mutant (Fig. 1A), suggesting that even this residual ROS production that could result from chloroplasts or differences in intracellular NADPH levels is under the control of RBOHs.

**Figure 1.**
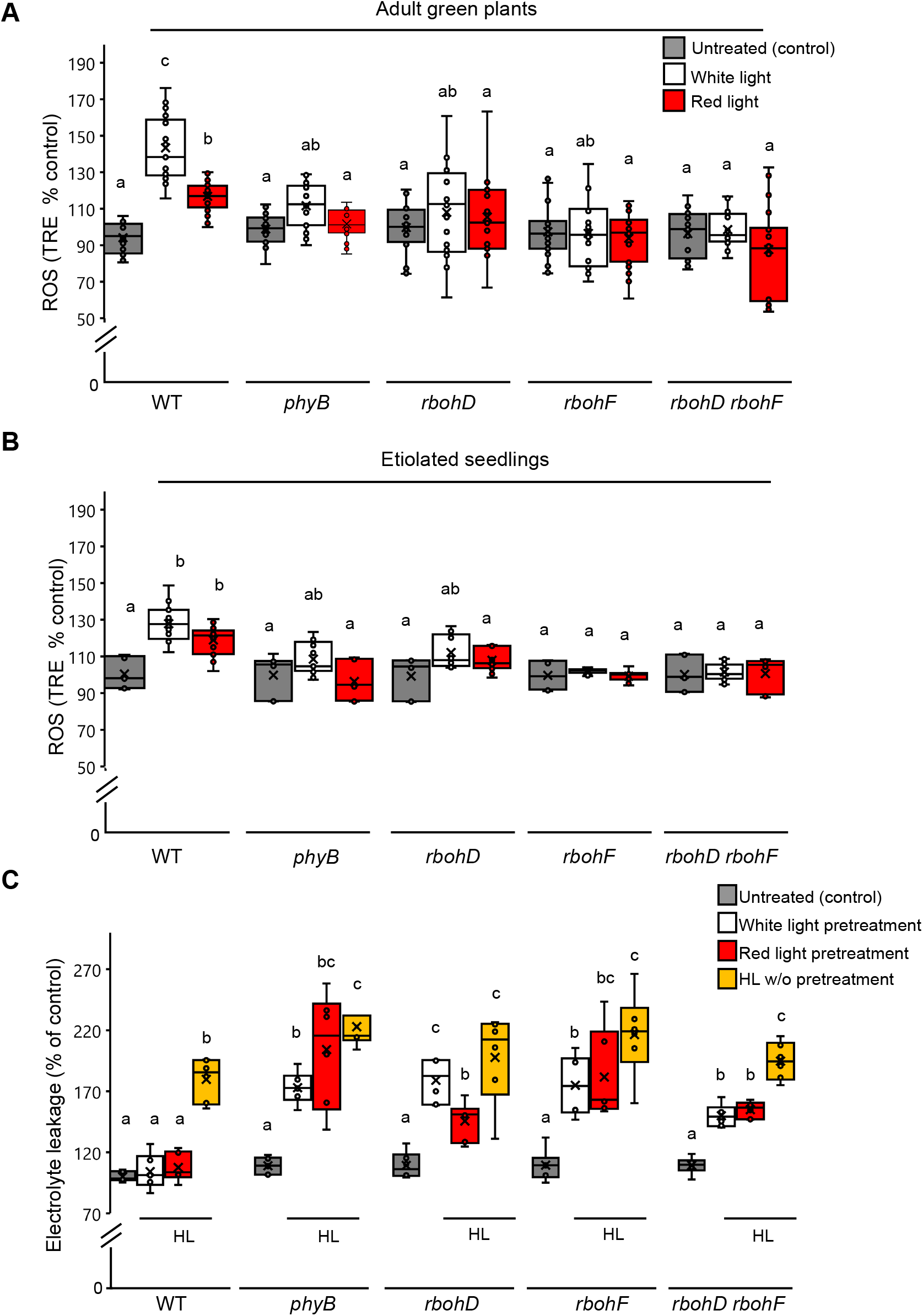
PhyB, RBOHD and RBOHF regulate ROS accumulation, physiological responses, and plant acclimation to excess light stress. (A) ROS accumulation in WT, *phyB*, *rbohD*, *rbohF*,and *rbohD rbohF* 4-week-old plants subjected to 10 min of excess white or red light. (B) Same as (A), but for 6-day-old etiolated seedlings grown on plates. (C) Acclimation of WT, *phyB*, *rbohD*, *rbohF*, and *rbohD rbohF* plants to excess light stress. Measurements are shown for untreated plants (control), plants that were pretreated with excess white or red light for 10 min, allowed to recover from 50 min and then subjected to excess white light for 45 min (pretreated), and plants that were treated with a 45 min excess white light without pretreatment (w/o pretreatment). Data is represented as percent of control (untreated plants) ± S.E. All experiments were repeated at least three times with 4 plants per biological replicate. N=12; Different letters denote statistical significance at P< 0.05 (ANOVA followed by a Tukey’s post hoc test). Abbreviations: DCF, 2’, 7’-Dichlorofluorescin; HL, highlight; phyB, phytochrome B; rbohD, NADPH/respiratory burst oxidase protein D; rbohF, NADPH/respiratory burst oxidase protein F; ROS, reactive oxygen species; TRE, total radiant efficiency; w/o, without; WT, wild-type.

To test the effect of phyB, RBOHD and RBOHF on physiological responses of plants to light stress, we measured the stomatal aperture closure response of plants to excess light stress (34–36). As shown in *SI Appendix* Fig. S1, treatment of mature green WT plants with excess white or red light for 10 min resulted in stomatal aperture closure. In contrast, treatment of *phyB*, *rbohD*, *rbohF*, or *rbohD rbohF* mutants did not. Because the absence of ROS accumulation (Fig. 1A) and stomatal responses (*SI Appendix* Fig. S1) could lead to an impaired acclimation of plants to excess light stress (34, 36, 37), we measured the acclimation of mature WT, *phyB*, *rbohD*, *rbohF*, and *rbohD rbohF* plants to a prolonged excess white light treatment following a short pretreatment with excess white or red light and an incubation period. As shown in Fig. 1C, pretreatment of WT plants with 10 min of excess white or red light, followed by an incubation of 50 min under controlled growth conditions, protected plants from a subsequent exposure to 45 min of excess white light (*i.e*., prevented leaf injury as measured by electrolyte leakage, compared to plants that were subjected to the 45 min excess light treatment without a 10 min pretreatment with excess white or red light). In contrast, pretreatment of *phyB*, *rbohD*, *rbohF*, or *rbohD rbohF* plants with excess white or red light failed to induce plant acclimation to a subsequent prolonged excess white light stress (Fig. 1C). The findings presented in Fig. 1 and *SI Appendix* Fig. S1 demonstrate that phyB, RBOHD and RBOHF are essential for ROS accumulation, physiological responses, and plant acclimation to excess light stress.

### PhyB, RBOHD and RBOHF regulate the expression of thousands of transcripts in response to excess light stress

To determine whether phyB, RBOHD and RBOHF function in the same signaling pathway that regulates ROS accumulation, physiological responses and plant acclimation to excess light stress (Fig. 1, *SI Appendix* Fig. S1), we subjected mature WT, *phyB*, *rbohD* and *rbohF* plants to a 10 min treatment with excess white (740 μmole photons s^−1^ m^−2^) or red (120 μmole photons s^−1^ m^−2^) light stress and studied their transcriptomics response using RNA-Seq analysis (Fig. 2; Datasets S1-S28). Quantitative RT-PCR (qPCR) analysis conducted on the different RNA samples obtained prior to the RNA-Seq analysis, revealed a complex response of the different mutants to the different treatments (*SI Appendix* Fig. S2). Some transcripts that were upregulated in WT but not *rbohD* or *rbohF* were still upregulated in *phyB* [*e.g., MYELOBLASTOSIS DOMAIN PROTEIN 30* (*MYB30*), *ZINC FINGER OF ARABIDOPSIS THALIANA 12* (*Zat12*) and *ASCORBATE PEROXIDASE 2* (*APX2*)], while others, such as *ZINC FINGER HOMEODOMAIN* 5 (ZHD5), that were upregulated in WT were suppressed in all mutants (*SI Appendix* Fig. S2). Venn diagrams comparing the transcripts significantly altered in WT and *phyB* in response to the excess white light treatment reveal that 3091 transcripts altered in WT by this treatment were not altered in the *phyB* mutant (Fig. 2A; Dataset S17). A similar comparison of transcripts altered in WT and the *rbohD* mutant revealed that 3460 transcripts altered in WT were not altered in *rbohD* (Fig. 2A; Dataset S18). Interestingly, an overlap of 2674 transcripts was found between the 3091 transcripts putatively regulated by phyB during light stress and the 3460 transcripts putatively regulated by RBOHD during light stress, suggesting that phyB and RBOHD co-regulate over 2500 transcripts during excess white light stress (Dataset S19). A similar comparison conducted between the response of *phyB* and *rbohF* revealed an overlap of 2446 transcripts (Fig. 2B; Dataset S20, S21). Interestingly, despite some differences in expression pattern between RBOHD and RBOHF in leaves (the latter being primarily expressed in vascular bundles; 32, 38), an overlap of 2217 transcripts was found between the transcripts common to phyB and RBOHD (2674) and phyB and RBOHF (2446), suggesting that phyB, RBOHD and RBOHF regulate a large proportion of the plant response to excess light stress (Fig. 2B; Dataset S22). A similar analysis conducted for the transcriptomics response of WT, *phyB*, *rbohF*, and *rbohD* to excess red light (*SI Appendix* Fig. S3; Datasets S23-S28) revealed an overlap of 353 transcript between the transcripts common to phyB and RBOHD (393) and phyB and RBOHF (490) in response to excess red light that should primarily activate phys (Fig. 2B; Dataset S28).

**Figure 2.**
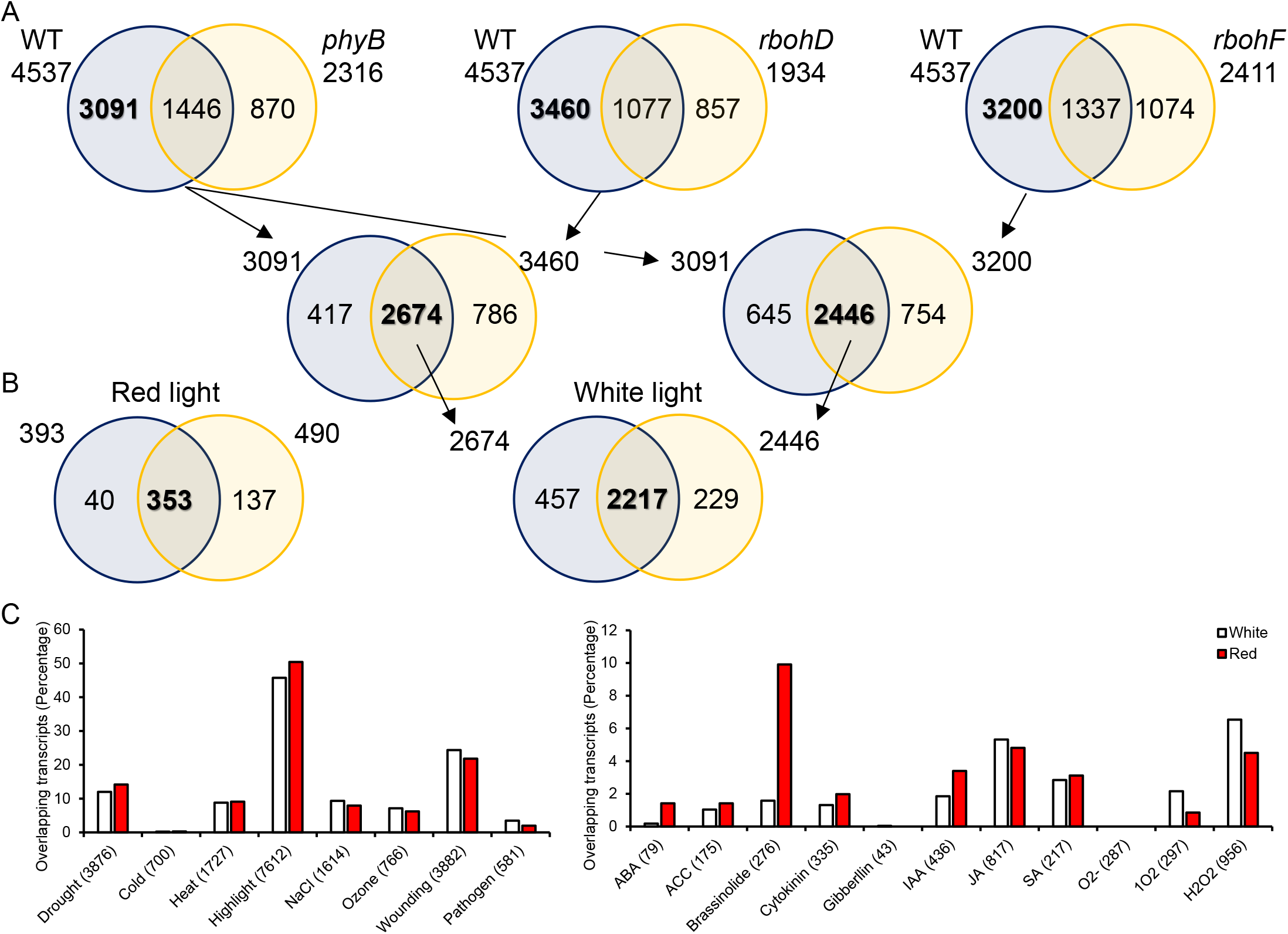
PhyB, RBOHD and RBOHF regulate the expression of thousands of transcripts in response to excess light stress. (A) Venn diagrams showing the overlap between transcripts altered in their expression in WT, *phyB*, *rbohD*, and *rbohF* plants in response to excess white light stress. (B) Venn diagram showing the overall between the transcripts altered in *phyB*, *rbohD*, and *rbohF* plants in response to excess red light stress. A complete analysis similar to (A),of the red light induced 353 transcripts that are co-regulated by phyB, RBOHD and RBOHF. Venn diagrams showing the overlap between transcripts altered in their expression in WT, *phyB*, *rbohD*, and *rbohF* plants in response to excess red light stress are shown in *SI Appendix* Fig. S3. (C) Precent representation of transcripts altered in response to different stresses, hormone treatments or ROS in the white (2217) and red (353) light response transcripts common to *phyB, rbohD*, and *rbohF* plants (from B and C). Abbreviations: ABA, abscisic acid; ACC, 1-aminocyclopropane 1-carboxylic acid; H_2_O_2_, hydrogen peroxide; IAA, indole-3-acetic acid; JA, jasmonic acid; O_2_^−^, superoxide;^1^O_2_, singlet oxygen; phyB, phytochrome B; rbohD, NADPH/respiratory burst oxidase protein D; rbohF, NADPH/respiratory burst oxidase protein F; ROS, reactive oxygen species; SA, salicylic acid; WT, wild-type.

An analysis of stress, hormone and ROS response transcripts found within these two groups of overlapping transcripts (2217 in response to white light, and 393 in response to red light), revealed that both of these groups contained a large number of high light-, wounding-, drought-, heat-, and salt-response transcripts (Fig. 2C). In addition, they contained a high number of H_2_O_2_-, brassinosteroid-, and jasmonic acid-response transcripts (Fig. 2C). These findings could suggest that similar to RBOHs, phyB is involved in the response of plants to many different stress conditions.

The results presented in Figs. 2 and *SI Appendix* Fig. S3 reveal that phyB, RBOHD and RBOHF co-regulate 1000s of transcripts during the response of plants to excess white light stress. Because RBOHD and RBOHF are thought to regulate plant responses via ROS production (29, 30, 39), and in the absence of phyB ROS does not accumulate in plants in response to excess light stress (red or white light; Fig. 1), it is likely that phyB functions upstream to RBOHD and RBOHF during plant responses to excess light stress.

### Complementing *phyB* with a cytosolic-restricted phyB protein recovers ROS production during excess light stress

The findings that phyB is required for RBOH-mediated ROS production during excess light stress (Fig. 1) and that phyB, RBOHD, and RBOHF co-regulate many transcripts during this response (Fig. 2), suggest that phyB regulates RBOH function. Although phyB was shown to interact with the PM and to exert some of its functions in the cytosol (40–42), the majority of phyB functions are mediated following its nuclear localization and interactions with various transcriptional regulators (43–49). To test whether phyB functions in the cytosol or nuclei to regulate RBOH function we expressed phyB fused in frame to glucocorticoid receptor 1 (GR1) in the *phyB* mutant (*phyB PHYB-GR1*). Previous studies showed that the phyB-GR1 hybrid protein is retained in the cytosol unless dexamethasone (DEX) is applied (50, 51). To validate the function of the phyB-GR1 system, we germinated WT, *phyB*, and *phyB PHYB-GR1* on plates in the presence or absence of DEX in the dark or under red light for 3 days. As shown in Fig. 3A, WT seedlings grown in darkness had an elongated hypocotyl. In response to red light however their hypocotyl was short. In contrast to WT, *phyB* seedlings did not respond to red light and had an elongated hypocotyl under dark or red-light conditions. Compared to WT or *phyB*, that were not responsive to the DEX treatment, the *phyB PHYB-GR1* line grown in the absence of DEX behaved like the *phyB* line, while the *phyB PHYB-GR1* grown in the presence of DEX behaved like the WT (Fig. 3A, *SI Appendix* S4A). These results confirm that in the absence of DEX the phyB-GR1 protein is retained in the cytosol, while in the presence of DEX it is mobilized to the nuclei (50, 51).

**Figure 3.**
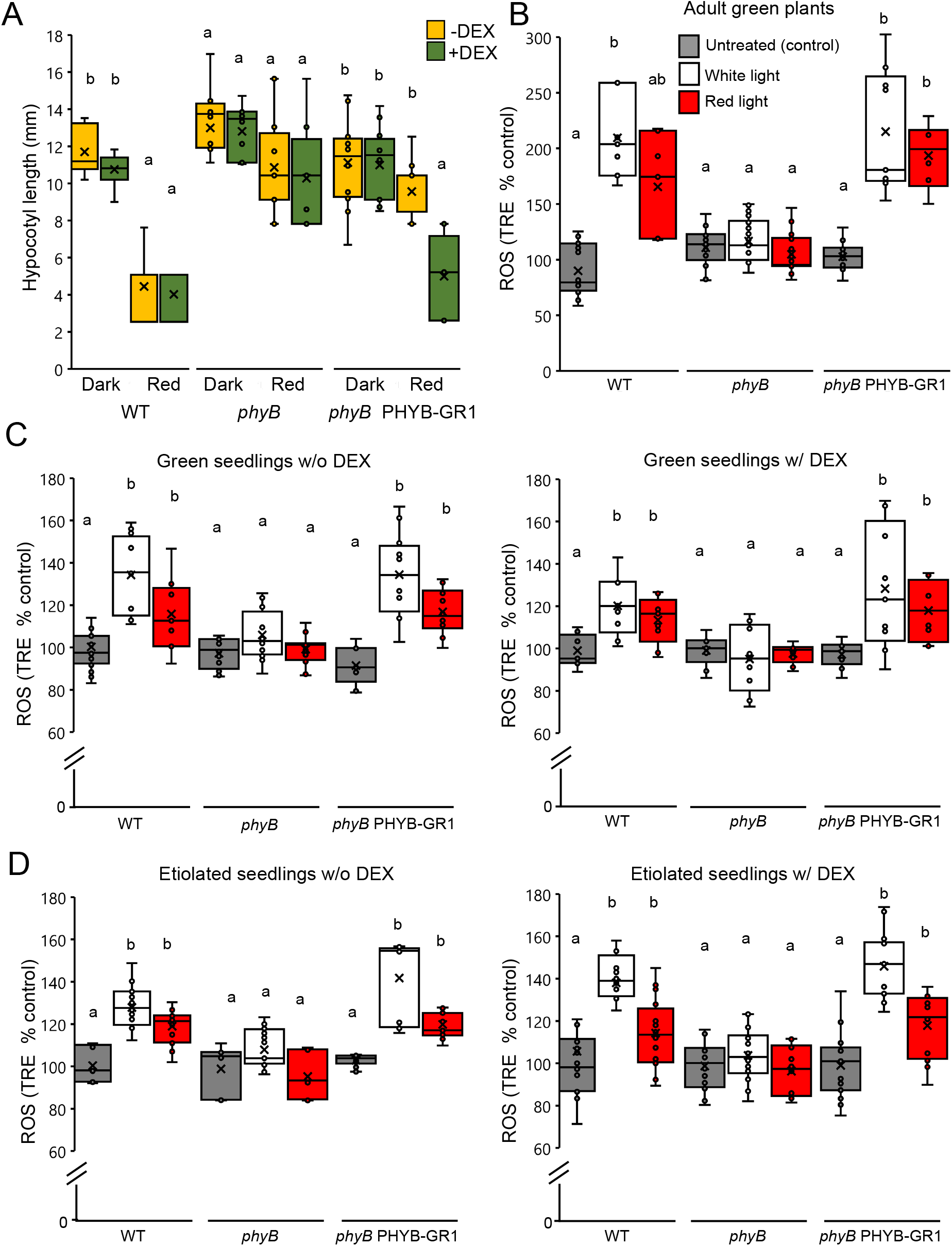
Complementing *phyB* with a cytosolic-restricted phyB protein recovers ROS production during excess light stress. (A) Hypocotyl length of WT, *phyB* and *phyB* PHYB-GR1 seedlings grown without DEX (yellow) or with DEX (green) in dark or under constant red light. (B) ROS accumulation in WT, *phyB*, and *phyB* PHYB-GR1 adult green plants subjected to 10 min of excess white or red light stress. (C) Same as (B), but for green seedlings grown on plates with or without DEX. (D) Same as (B), but for etiolated seedlings grown on plates with or without DEX. Data is represented as percent of control (untreated plants) ± S.E. Different letters denote statistical significance at P< 0.05 (ANOVA followed by a Tukey’s post hoc test). Abbreviations: DEX, dexamethasone; GR1, Glucocorticoid Receptor; HL, highlight; phyB, phytochrome B; ROS, reactive oxygen species; TRE, total radiant efficiency; w/o, without; WT, wild-type.

To test whether phyB functions in the cytosol or nuclei to regulate ROS accumulation during excess light stress, we subjected mature WT, *phyB*, and *phyB PHYB-GR1* plants (*SI Appendix* Fig. S4B) to excess white or red light stress in the absence of DEX (hybrid protein is restricted to the cytosol) and measured ROS accumulation. As shown in Fig. 3B, excess white or red light caused enhanced ROS accumulation in WT or *phyB PHYB-GR1* plants, suggesting that the phyB-GR1 protein present in the cytosol is sufficient to trigger ROS production under conditions of excess light stress. To test the effect of DEX on ROS production in WT, *phyB* and *phyB PHYB-GR1* plants, we grew seedlings of these lines in the presence or absence of DEX and treated them with excess white or red light. As shown in Fig. 3C, the presence of DEX did not prevent the accumulation of ROS in WT or *phyB PHYB-GR1* plants, suggesting that not all the phyB-GR1 protein localized to the nuclei in *phyB PHYB-GR1* plants following DEX application, or that the phyB-GR1 protein can also induce ROS formation upon localization to the nuclei. Similar results were found with etiolated seedlings of the different lines grown in the presence or absence of DEX and subjected to excess white or red light (Fig. 3D). The results presented in Fig. 3 demonstrate that phyB does not need to localize into the nuclei to trigger ROS production in response to excess light stress.

### Plant acclimation to excess light stress requires nuclear localization of phyB

The findings that phyB can regulate ROS production during excess light stress without entering the nuclei (Fig. 3) prompted us to test whether it can also trigger plant acclimation to excess light stress while being restricted to the cytosol. For this purpose, we developed a new assay to measure the acclimation of WT, *phyB* and *phyB PHYB-GR1* seedlings growing in liquid media in the presence or absence of DEX to excess light stress. As shown in Fig. 4, WT seedlings could acclimate to a prolonged excess white light stress treatment following a pretreatment with a short excess white or red light stress followed by incubation, regardless of the presence or absence of DEX. In contrast, and regardless of the presence or absence of DEX, *phyB* seedlings were unable to acclimate the excess light stress treatment (Fig. 4). Interestingly, while *phyB PHYB-GR1* seedlings were unable to acclimate to the excess white light treatment in the absence of DEX, they were able to partially acclimate to it in the presence of DEX. This finding suggests that nuclear localization of phyB could be required for plant acclimation to excess light stress.

**Figure 4.**
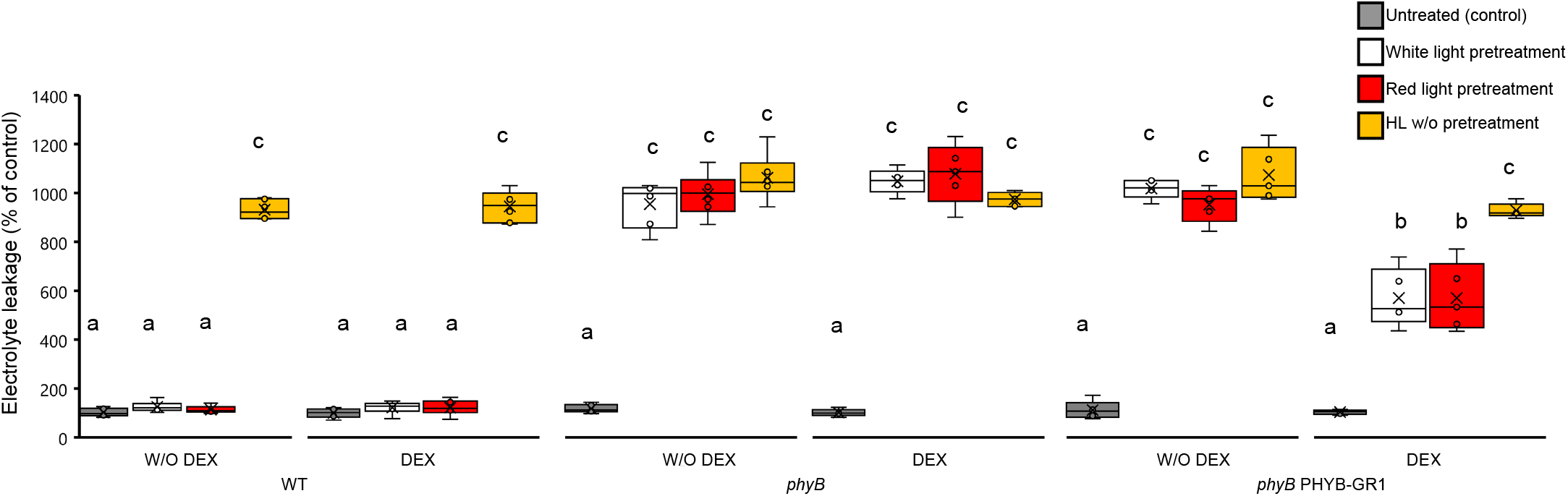
Plant acclimation to excess light stress requires nuclear localization of phyB. Acclimation of WT, *phyB* and *phyB* PHYB-GR1 seedlings grown in liquid in the presence or absence of DEX to excess white light stress. Measurements are shown for untreated seedlings (control), seedlings that were pretreated with excess white or red light for 10 min, allowed to recover from 50 min and then subjected to excess white light for 45 min (pretreated), and seedlings that were treated with a 45 min excess white light without pretreatment (w/o pretreatment). Data is represented as percent of control (untreated plants) ± S.E. All experiments were repeated at least three times with 45 seedlings per biological replicate. Different letters denote statistical significance at P< 0.05 (ANOVA followed by a Tukey’s post hoc test). Abbreviations: DEX, dexamethasone; GR1, Glucocorticoid Receptor; HL, highlight; phyB, phytochrome B; ROS, reactive oxygen species; TRE, total radiant efficiency; w/o, without; WT, wild-type.

### PhyB is required for ROS production in response to wounding, heat, cold, and pathogen infection

We previously reported that RBOHD is required for rapid ROS production and systemic signaling in *Arabidopsis* in response to many different abiotic and biotic stimuli (33, 34, 52). The findings that phyB is required for rapid ROS production in response to excess light stress (Fig. 1), that phyB and RBOHs are required for the expression of the same set of transcripts during light stress (Fig. 2), and that phyB can regulate rapid ROS production even if it is restricted to the cytosol (Fig. 3), prompted us to test whether phyB is required for rapid ROS production and systemic signaling during the response of *Arabidopsis* to other stresses. As shown in Fig. 5A-D, phyB is required for rapid local and systemic ROS accumulation in response to excess light, cold, heat, or pathogen infection. In contrast, phyB is required for systemic, but not local, ROS accumulation during wounding (Fig. 5E). The findings presented in Fig. 5 suggest that phyB is required for the rapid ROS accumulation response of plants to several different abiotic and biotic stresses, potentially functioning as part of a signaling module with RBOHD and/or RBOHF.

**Figure 5.**
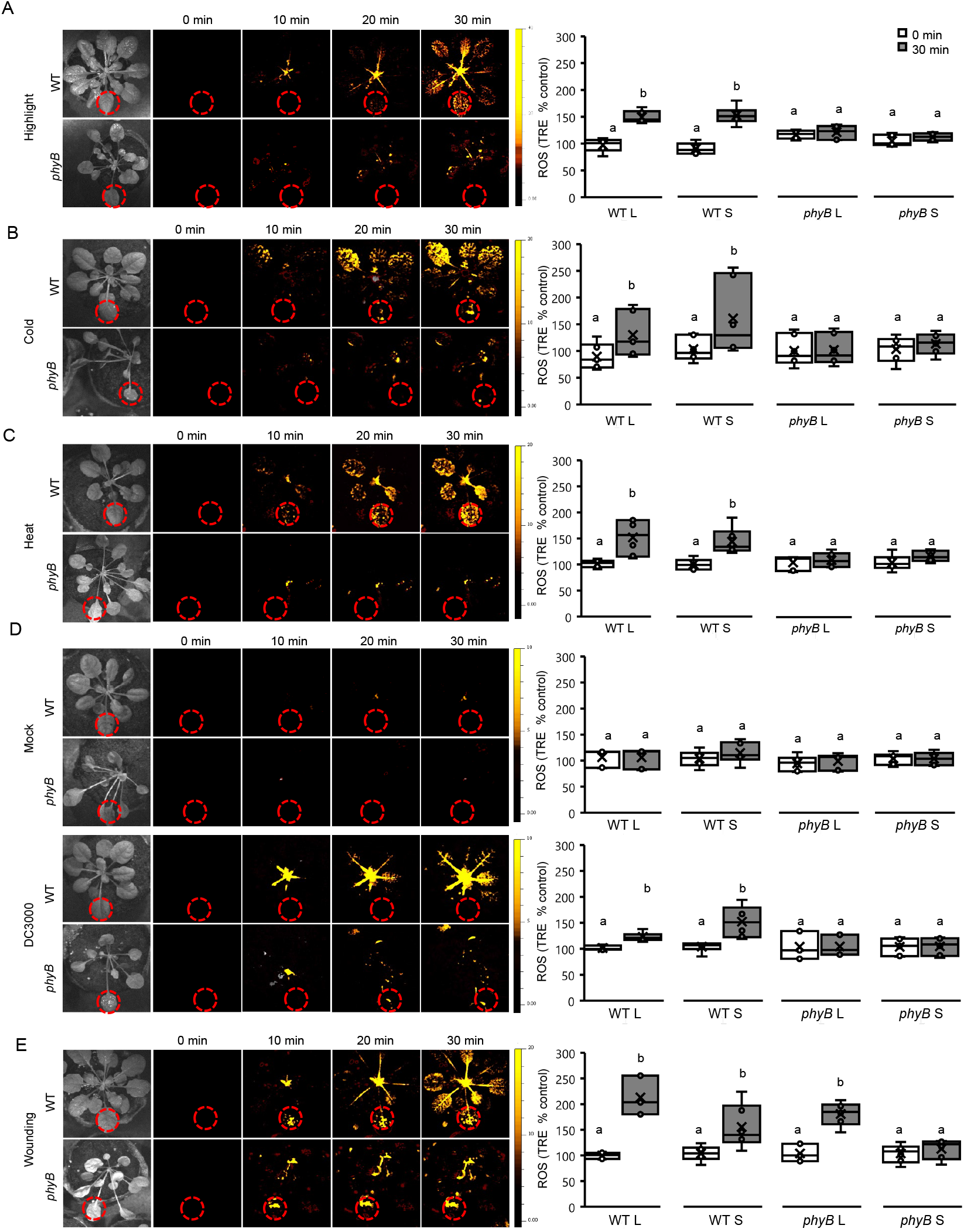
PhyB is required for ROS production in response to wounding, heat, cold, and pathogen infection. (A) Representative time lapse images (left) and quantitative bar graphs (right) of ROS accumulation in local (L) and systemic (S) leaves (local leaf is marked with red broken line circle) of WT and *phyB* plants subjected to an excess light stress treatment applied to the local leaf. (B) same as (A), but for cold stress applied to the L leaf. (C) same as (A), but for heat stress applied to the L leaf. (D) same as (A), but for mock or *P. syringae* DC3000 infection applied to the L leaf. (E) same as (A), but for mechanical wounding applied to the L leaf. Data is represented as percent of control (0 min) ± S.E. Different letters denote statistical significance at P< 0.05 (ANOVA followed by a Tukey’s post hoc test). Abbreviations: phyB, phytochrome B; L, local; ROS, reactive oxygen species; S, systemic; TRE, total radiant efficiency; WT, wild-type.

### PhyB is required for ROS accumulation during excess light stress in rice

We previously reported that RBOHA is required for ROS production during excess light stress in rice (25). To test whether phyB is also involved in the response of rice to excess light stress, we subjected WT, *phyB*, *rbohA*, and *rbohB* rice plants to excess white or red light stress and measured ROS accumulation using our whole plant live ROS imaging method (33). As shown in Fig. 6, WT rice plants subjected to excess light stress accumulated ROS. In contrast, rice *phyB*, *rbohA*, and *rbohB* mutants, did not. These findings demonstrate that phyB could play an important role in ROS production during excess light stress in other plants.

**Figure 6.**
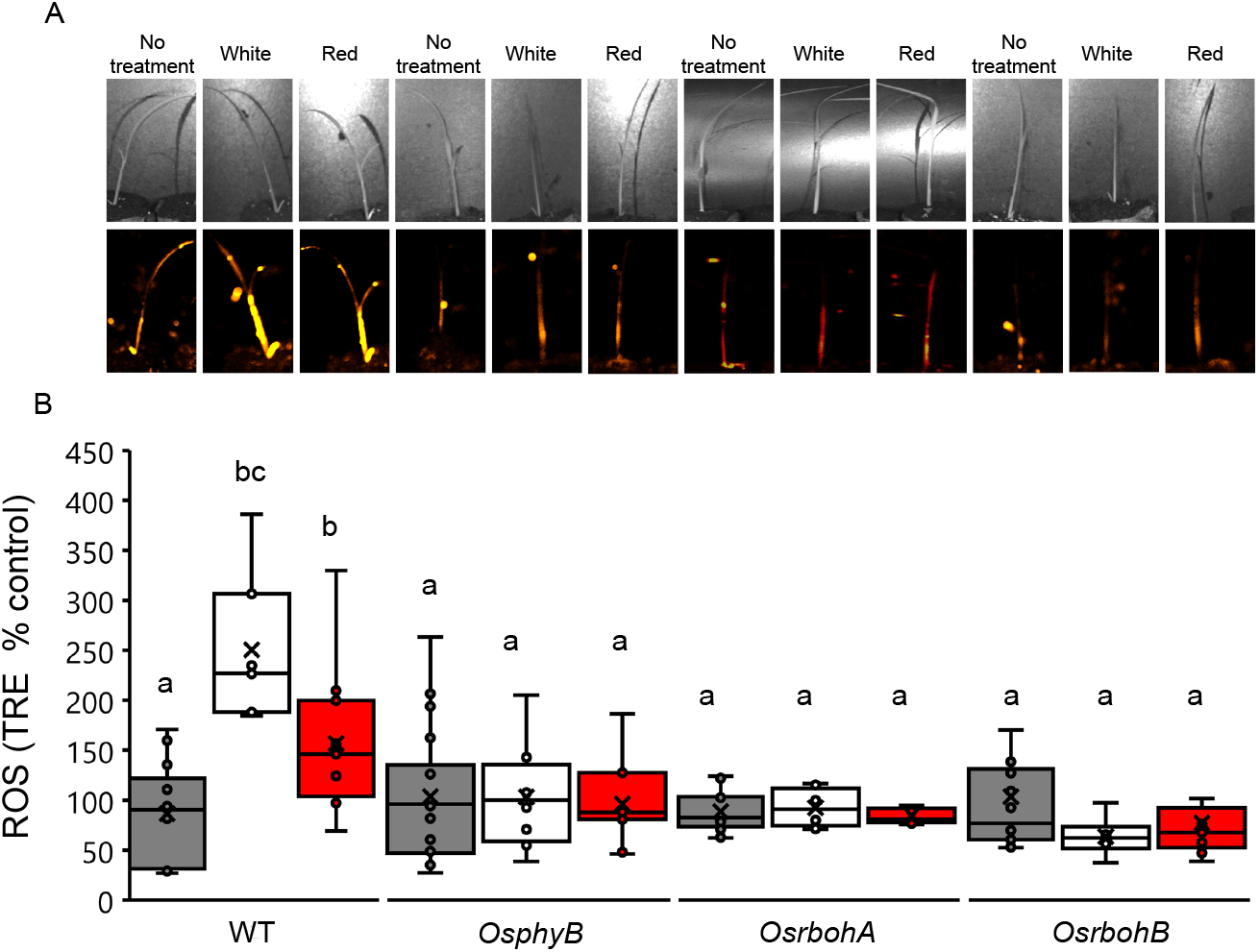
PhyB is required for ROS accumulation during excess light stress in rice. (A) representative images of whole-plant ROS accumulation in WT, *OsphyB, OsrbohA*, and *OsrbohB* rice plants subjected to 10 min of excess white or red light. (B) Quantitative analysis of ROS accumulation in WT, *OsphyB, OsrbohA*, and *OsrbohB* rice plants subjected to 10 min of excess white or red light. All experiments were repeated at least three times with 10 plants per biological replicate. Data is represented as percent of control (untreated plants) ± S.E. Different letters denote statistical significance at P< 0.05 (ANOVA followed by a Tukey’s post hoc test). Abbreviations: ROS, reactive oxygen species; DCF, 2’, 7’-Dichlorofluorescin; WT, wild-type; phyB, phytochrome B; rbohD, NADPH/respiratory burst oxidase protein D; rbohF, NADPH/respiratory burst oxidase protein F; TRE, total radiant efficiency.

## Discussion

The photoreceptor protein phyB plays a key role in the sensing of light and heat leading to the activation and/or modulation of many different developmental programs and environmental responses (1, 2). Recent studies revealed that in addition to light and heat, phyB is involved in responses of plants to chilling, salt, drought, pathogen, and cold stresses (43, 46, 53–55). To regulate plant responses to environmental stimuli, phyB interacts with, or affects the function of, multiple proteins including transcriptional regulators such as PHYTOCHROME-INTERACTING TRANSCRIPTION FACTORS (PIFs), C-REPEAT/DRE-BINDING FACTORS (CBFs), MYELOBLASTOSIS DOMAIN PROTEINS (MYBs), and CALMODULIN-BINDING TRANSCRIPTION ACTIVATOR (CAMTA), binds to different promoters, and/or associates with large multi protein complexes called photobodies (1–6, 43–49). PhyB primarily functions therefore in the nuclei to alter gene expression. Here we reveal that in addition to its nuclear functions, phyB acts in the cytosol to regulate ROS production by RBOHs during excess light stress (Fig. 3). Moreover, we show that phyB is essential for ROS production during excess light stress in both the dicot Arabidopsis (Fig. 1) and the monocot rice (Fig. 6), suggesting that its function upstream to RBOHs is conserved and widespread in plants. We further show that phyB is required for local and/or systemic ROS accumulation in plants in response to pathogen infection, cold, heat, or wounding (Fig. 5). These findings shed new light on phyB function in plants, *i.e*., the regulation of ROS production at the apoplast in response to multiple stimuli. This new role is highly important for our understanding of plant biology since ROS production is central for plant responses to many different environmental conditions and stresses (14, 15, 17, 24, 56). PhyB could therefore function in both the nuclei and the cytosol to coordinate apoplastic ROS production during stress with different transcriptional responses (Fig. 7). Supporting this hypothesis are our findings that although the cytosolic localization of phyB was sufficient to activate ROS production (Fig. 3), it was not sufficient to induce plant acclimation (Fig. 4). Instead, nuclear functions of phyB appear necessary to induce plant acclimation to excess light stress (Fig. 4).

**Figure 7.**
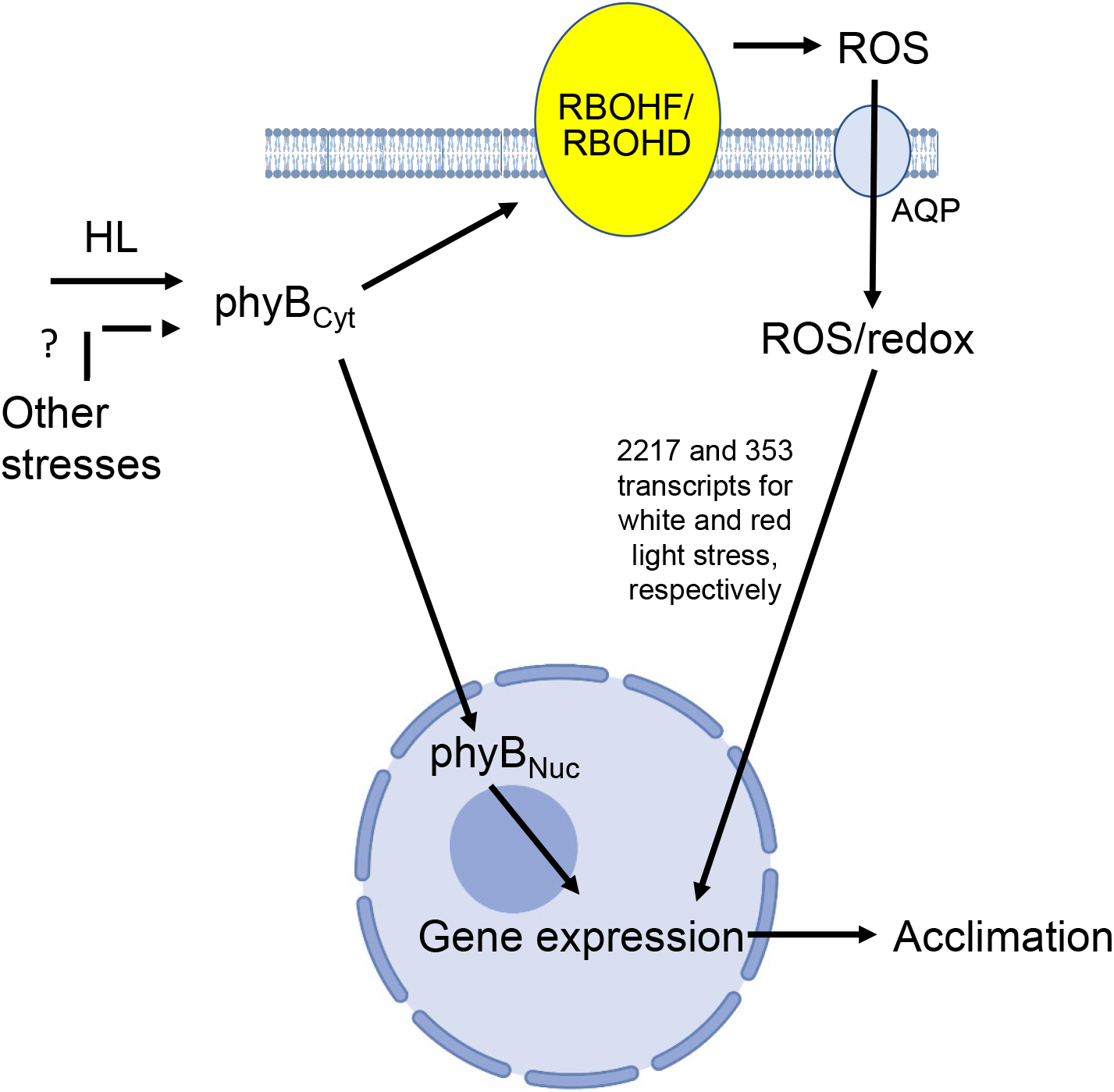
Model for phyB-RBOH signaling during the response of plant to stress. PhyB is shown to integrate different environmental stresses and to activate ROS production by RBOHD and RBOHF at the apoplast. PhyB can mediate this function even if it is restricted to the cytosol. To induce plant acclimation to stress, phyB needs however to localize to the nucleus. In response to excess white light stress, pyhB, RBOHD and RBOHF are shown to regulate the expression of 1000s of transcripts that depend on ROS production at the apoplast. ROS at the apoplast is thought to enter into the cytosol via aquaporins. Abbreviations: AQP, aquaporin; Cyt, cytosolic; HL, highlight; phyB, phytochrome B; Nuc, nuclear; PM, plasma membrane; rbohD, NADPH/respiratory burst oxidase protein D; rbohF, NADPH/respiratory burst oxidase protein F; ROS, reactive oxygen species.

Our findings that in the absence of phyB ROS do not accumulate in plant cells in response to excess light stress (Figs. 1, 6), that phyB can regulate ROS production without localizing to the nucleus (Fig. 3), and that phyB function overlaps with that of RBOHs in regulating transcriptomics responses during excess light stress (Fig. 2), strongly suggests that phyB functions upstream to RBOHs (Fig. 7). Taking into consideration the broad signaling function of RBOHs in mediating ROS production during biotic and abiotic stresses (29, 30, 39), and the importance of phyB for many of these responses (43–49), it is likely that a phyB-RBOH regulatory module is involved in ROS production during plant responses to many different stresses. Several mechanisms could link phyB function in the cytosol with the PM localized RBOHs. PhyB was shown, for example, to associate with the plant PM and could directly bind to or interact with RBOHs (40). In addition, phyB contains a Histidine kinase domain and RBOHs are regulated by phosphorylation (3–7, 29, 30, 39, 41, 42). PhyB was also found to interact with CALCIUM DEPENDENT PROTEIN KINASE 1 (CPK1) that phosphorylates CBL-INTERACTING SERINE/THREONINE-PROTEIN KINASE 1 (CIPK1) which phosphorylates CALCINEURIN B-LIKE PROTEIN 9 (CBL9) and CBL-INTERACTING SERINE/THREONINE-PROTEIN KINASE 26 (CIPK26), a known activator of RBOHs (https://thebiogrid.org/; 57, 58). These possible regulatory mechanisms should be examined in future studies.

The possible broad function of phyB in regulating ROS production during plant responses to different stresses is supported by previous studies showing the involvement of phyB in plant responses to some of these stresses (43, 46), by our findings of a large overlap in transcripts regulated by phyB and RBOHs (Fig. 2A), and by the nature of the transcripts regulated by phyB and RBOHs (Fig. 2C). These include a high proportion of excess light-, drought-, heat-, salinity-, pathogen-, wounding-, and ozone-response transcripts. In addition, they contain H_2_O_2_-, jasmonic acid-, salicylic acid-, and auxin-response transcripts (Fig. 2C). Taken together, these data suggest that the phyB-RBOH signaling module may play a central role in regulating ROS levels and plant acclimation to many different abiotic and biotic stresses in plants (Fig. 7).

Our findings that the majority of ROS accumulating in plant cells during excess light stress is dependent on RBOH function and not produced by chloroplasts, peroxisomes or mitochondria are intriguing (Figs. 1, 6; 26, 31). Especially since ROS were found to be produced in these organelles and to have an important regulatory function (14–17). One possible explanation to these findings is that ROS produced in different organelles during excess light stress trigger retrograde signaling pathways that affect nuclear responses but are simultaneously scavenged within these organelles and do not accumulate to high levels. It is also possible that low levels of ROS produced in these organelles diffuse into the cytosol to regulate redox and signaling pathways, but that these levels of ROS are too low for our method to detect. Because ROS can accumulate in the apoplast to high levels without having a toxic effect (26, 33, 34, 52), it is also possible that apoplastic ROS production by RBOHs could serve as a transient stress memory mechanism, as previously shown for light stress in *Arabidopsis* that triggered an RBOHD-dependent 3-6 hour high apoplastic ROS production state (26). During stress, ROS could therefore be produced in organelles to trigger different retrograde signaling pathway, as well as in the apoplast to serve as a ROS reservoir (much like calcium is stored in the apoplast and other compartments; 59), and a delicate interplay between retrograde mechanisms and low ROS levels that enter the cytosol from different organelles or the apoplast could control different signaling pathways and lead to acclimation. In this respect, it should be mentioned that at least one retrograde pathway involving the signaling metabolite methylerythritol cyclodiphosphate (MEcPP) and the transcription factor CAMTA3 was recently shown to regulate phyB function (44). Light stress could therefore be sensed by phyB, or chloroplasts, which could affect phyB function via retrograde signaling, and alter transcriptional responses. Further studies are of course needed to determine how different signals generated in different compartments are coordinated to trigger plant acclimation to excess light or other stresses.

In addition to understanding the mode of coordination between different compartments during stress, attention should also be given to the temporal role ROS play in plant responses (57). In the current study we used a 10 min excess light stress treatment. This time point could represent an early response time point for excess light stress in which apoplastic ROS accumulation is predominant. In later or earlier time points, or following chronic exposure to excess light stress, other compartments, such as chloroplasts or peroxisomes, could also contribute to ROS production. Further studies are therefore needed to determine the different temporal and spatial modes of ROS production and signaling during excess light stress and the role(s) that phyB plays in these responses. With respect to the extent of plant exposure to excess light stress used in the current study (10 min) it is important to note that the exposure of plants to this amount of excess light stress is sufficient to induce plant acclimation (Fig. 4; 34, 60, 61). We are therefore relatively confident that the role of ROS and phyB identified by the current study is biologically significant.

## Materials and Methods

### Plant material and growth conditions

Wild-type *Arabidopsis thaliana* (cv Columbia), *rbohD*(AT5G47910; 62), *rbohF* (AT1G64060; 62), *rbohD rbohF* (63),*phyB* (AT2G18790; 64) and *phyB PHYB-GR1* (51) were tested in this study. Four-week-old plants were grown on peat pellets (Jiffy-7; Jiffy International, Kristiansand, Norway), at 21°C, under 10hr/14hr light/dark (50 μmole photons s^−1^ m^−2^) conditions, and used for ROS quantification, acclimation, stomatal aperture, and RNA analysis. To study green or etiolated seedlings, seeds from the above genotypes were germinated and grown on 0.5 X Murashige and Skoog media (Caisson Labs, Smithfield, UT, USA) for 7 days, before light stress and ROS quantification. Plates were supplemented with 0.1 mM dexamethasone (Sigma-Aldrich, St. Louis, MO, USA) for phyB PHYB-GR1 experiments. For acclimation experiments with and without dexamethasone treatments, seedlings were grown in liquid cultures of 0.25 X Murashige and Skoog in 6-well-plates. Rice (*Oryza sativa* L.) plants studied included Nipponbare WT, *OsphyB, OsrbohA-1* and <u>*OsrbohB*</u> (25). Rice plants were grown on peat pellet for 3 weeks in 16hr/8hr light/dark (100 μmole photons s^−1^ m^−2^) conditions before stress and ROS quantification.

### Stress treatments

Whole plants were subjected to light stress for 10 or 50 minutes. White light stress (740 μmole photons s^−1^ m^−2^; *SI Appendix* Fig. S5A) was applied using an LED array (BESTVA, Commerce, CA, USA) and covered a spectrum of 300-800 nm, while red-light stress (120 μmole photons s^−1^ m^−2^; *SI Appendix* Fig. S5B) was applied using an LED array (DUOSTRIP I033 light diodes array, LEDdynamics, Randolph, VT, USA) with a spectrum of 600-700 nm in a dark room. ROS measurements of systemic stress signaling were performed as previously described (33, 34), following a 2 min local light stress treatment (1700 μmole photons s^−1^ m^−2^) generated by a ColdVision fiber optic LED light source (Schott, Mainz, Germany). Wounding was applied by simultaneously injuring a single leaf with 30 dresser-pines (33), cold stress was applied by placing an ice cube on a single leaf for 2 min (49), heat stress was induced by placing a heat block 2 cm away from the treated leaf (34) and following leaf temperature using an infra-red camera (FLIR Wilsonville,OR, USA), and pathogen attack was applied by dipping a single leaf in a tube with Pseudomonas syringae DC3000 in phosphate buffer (for mock treatment, the leaf was dipped in a tube with the buffer without the bacteria; 33).

### Whole plant live ROS imaging, plant acclimation and stomatal aperture measurements

ROS accumulation was measured by enhanced fluorescence of oxidized DCF as previously described (33). Plants were fumigated with H2DCFDA solution [50uM H2DCFDA (Sigma-aldrich, St. Louis, MO, USA), 0.001% Silwet L-77 (Sigma-aldrich, St. Louis, MO, USA) in 50 mM phosphate buffer pH7.4] for 30 min (33). Internalized and oxidized DCF fluorescence (Ex./Em. 480nm/520nm) was measured using the IVIS Lumina S5 platform (PerkinElmer, Waltham, MA, USA). Accumulation of oxidized DCF was calculated using the IVIS Living Image 4.7.2 software (PerkinElmer, Waltham, MA, USA). Acclimation was performed as described previously (33, 34) with some modifications. Whole rosettes of Arabidopsis plants were pretreated with white or red light for 10 min. Following the pretreatment, plants were incubated in ambient light for 50 min. Following the recovery period, plants were put under high light (740 μmole photons s^−1^ m^−2^, 300-800 nm) for 45 min. Four leaves were collected and submerged in deionized H_2_O for 60 min in a tube on a rotator. The solution conductivity was measured with a conductivity meter (Okaton CON 510; Cole-Parmer, Vernon Hills, IL, USA) and then the tubes with the leaves were boiled for 20 min. Following cooling down to room temperature, the solution conductivity was measured again. The percentage of conductivity of the fresh sample from the boiled samples was used to calculate electrolyte leakage, compared to samples from untreated plants (without high light and pretreatment). For control, conductivity from untreated plants (without high light and pretreatment) and high light with pretreatment plants was measured as well. Acclimation of DEX treated and untreated seedlings was performed with 1-week-old seedlings grown in liquid 0.25 X Murashige and Skoog media (Caisson Labs, Smithfield, UT, USA). Seedlings were subjected to the same light treatments as described above, and conductivity was measured in the liquid growth media. Stomatal aperture measurements were performed as described previously (26).

### RNA extraction and transcript expression analysis

A total of 60 leaves were pooled from 15 different plants for each biological repeat (3 biological repeats were used for each time point and genotype) of WT, *phyB, rbohD* or *rbohF* subjected to 0 or 10 min of white light or red light, and frozen in liquid nitrogen before grinding and isolation of RNA using RNeasy Plant mini kit (Qiagen, Hilden, Germany). Isolated RNA was used as template for cDNA synthesis using PrimeScript RT Reagent Kit (Takara Bio, Kusatsu, Japan). Transcript expression was quantified by qRT-PCR using iQ SYBR Green supermix (Bio-Rad Laboratories, Hercules, CA, USA) using the specific primers in *SI Appendix* Dataset S29. RNA libraries for sequencing were prepared using standard Illumina protocols and RNA sequencing was performed by Novogene (Sacramento, CA, USA) using NovaSeq 6000. Read quality control was performed using FastQC v1.20.0 (https://www.bioinformatics.babraham.ac.uk/projects/fastqc/), followed by alignment of reads onto the Arabidopsis reference genome (The Arabidopsis Information Resource [TAIR] 65; https://www.arabidopsis.org/) using STAR aligner v2.4.0.1 (https://github.com/alexdobin/STAR) and analysis of differential gene expression using DESeq2 v1.20.0 (34, 66; https://bioconductor.org/packages/release/bioc/html/DESeq2.html). The genome index was built using the gene annotation file (gff file) downloaded from TAIR 10. Differences in expression were quantified as the logarithm of the ratio of mean normalized counts between two conditions (log fold change). Differentially abundant transcripts were defined as those that have a log fold change with an adjusted P < 0.05 (negative binomial Wald test followed by a Benjamini-Hochberg correction, both integral to the DESeq2 package). Differentially expressed genes were classified into upregulated or downregulated based on significant positive or negative log fold change values, respectively. Venn diagram overlaps were calculated using VIB Bioinformatics and Evolutionary Genomics web tool (http://bioinformatics.psb.ugent.be/webtools/Venn/). Raw and processed RNA-seq data files were deposited in the Gene Expression Omnibus (https://www.ncbi.nlm.nih.gov/geo/) under the accession number GSE188732.

### Statistical analysis

All experiments were repeated at least three times with at least 3 biological repeats. Box plots graphs are presented with ± SE. ANOVA was followed by a Tukey’s post hoc test. Different letters denote statistical significance at p < 0.05.

## Acknowledgments

We thank Dr. Enamul Huq for the *phyB PHYB-GR1* seeds. This work was supported by funding from the National Science Foundation (IOS-2110017, IOS-1353886, MCB-1936590, IOS-1932639), Interdisciplinary Plant Group and The University of Missouri.

## Figures Legends

**Figure S1.**
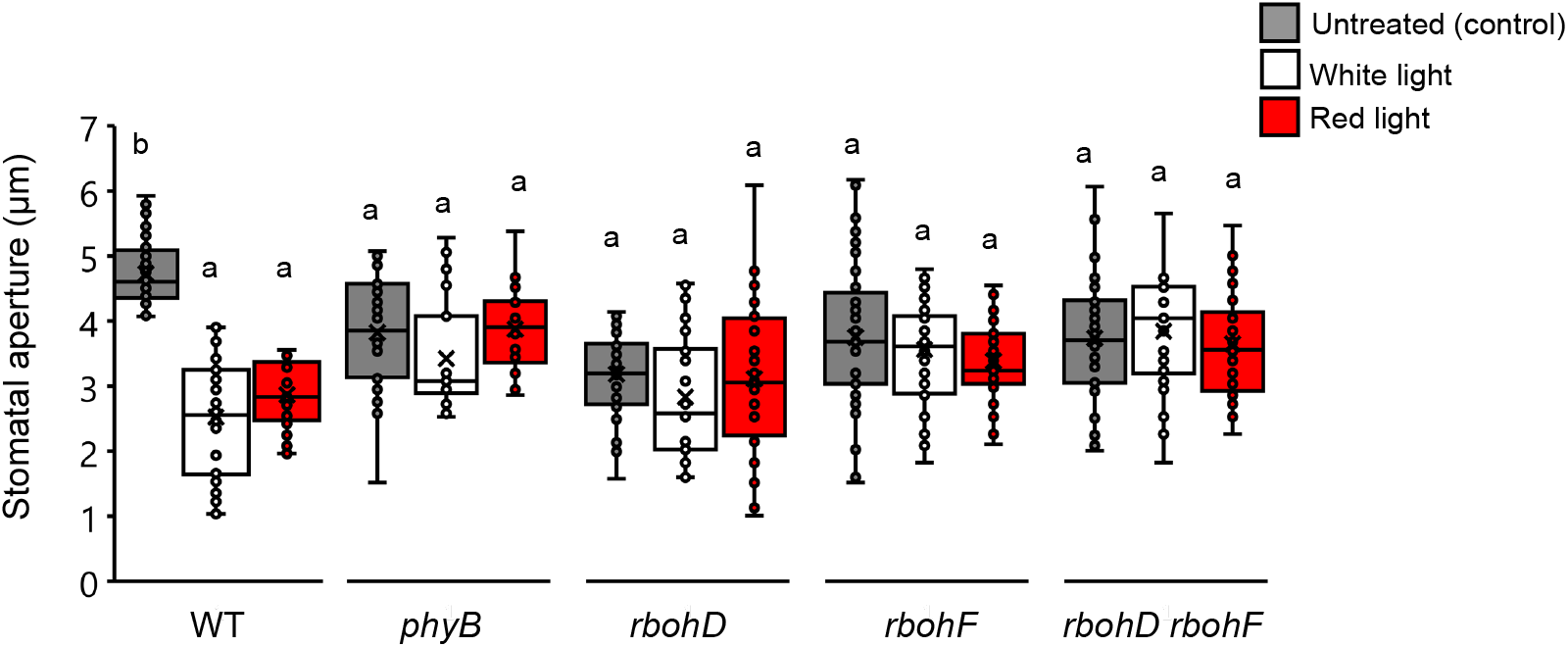
Stomatal aperture of WT, *phyB, rbohD, rbohF*, and *rbohD rbohF* plants subjected to a 10 min of excess white light or red-light treatment. All experiments were repeated at least three times with 4 plants per biological replicate. ANOVA followed by a Tukey’s post hoc test; N=12, P-value<0.05. Abbreviations: HL, highlight; phyB, phytochrome B; rbohD, respiratory burst oxidase homolog D; rbohF, respiratory burst oxidase homolog F; WT, wild-type.

**Figure S2.**
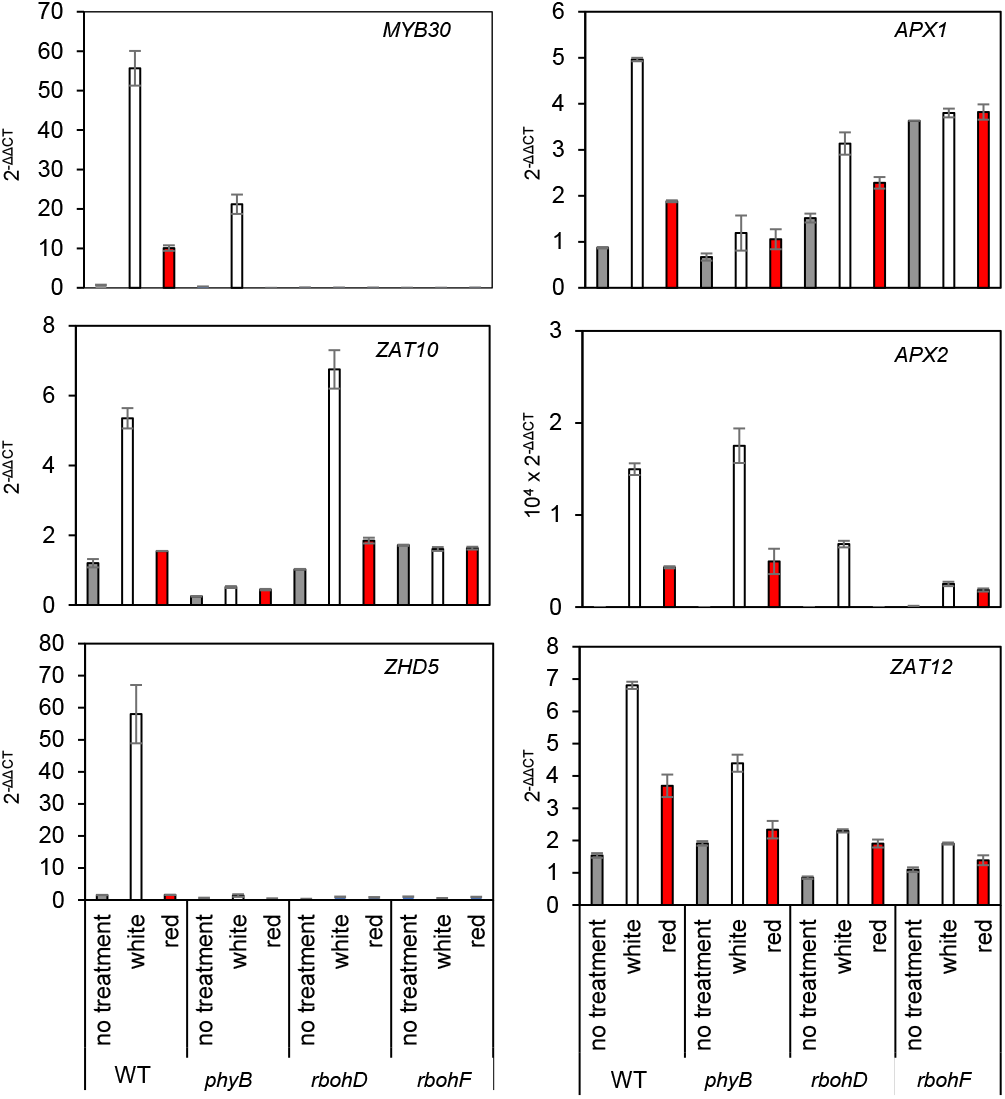
Transcript expression (qPCR) analysis following excess white or red light treatment of WT, *phyB, rbohD*, and *rbohF* plants. Results were calculated as 2^-ΔΔCT^ normalized with internal control and untreated samples. Abbreviations: APX1, ascorbate peroxidase 1; APX2, ascorbate peroxidase 2; HL, highlight; MYB30, myeloblastosis domain protein 30; PCR, polymerase chain reaction; phyB, phytochrome B; rbohD, respiratory burst oxidase homolog D; rbohF, respiratory burst oxidase homolog F; ROS, reactive oxygen species; WT, wildtype; ZAT10, Zinc finger of *Arabidopsis thaliana* 10; ZAT12, Zinc finger of Arabidopsis thaliana 12; ZHD5, Zinc finger homeodomain 5.

**Figure S3.**
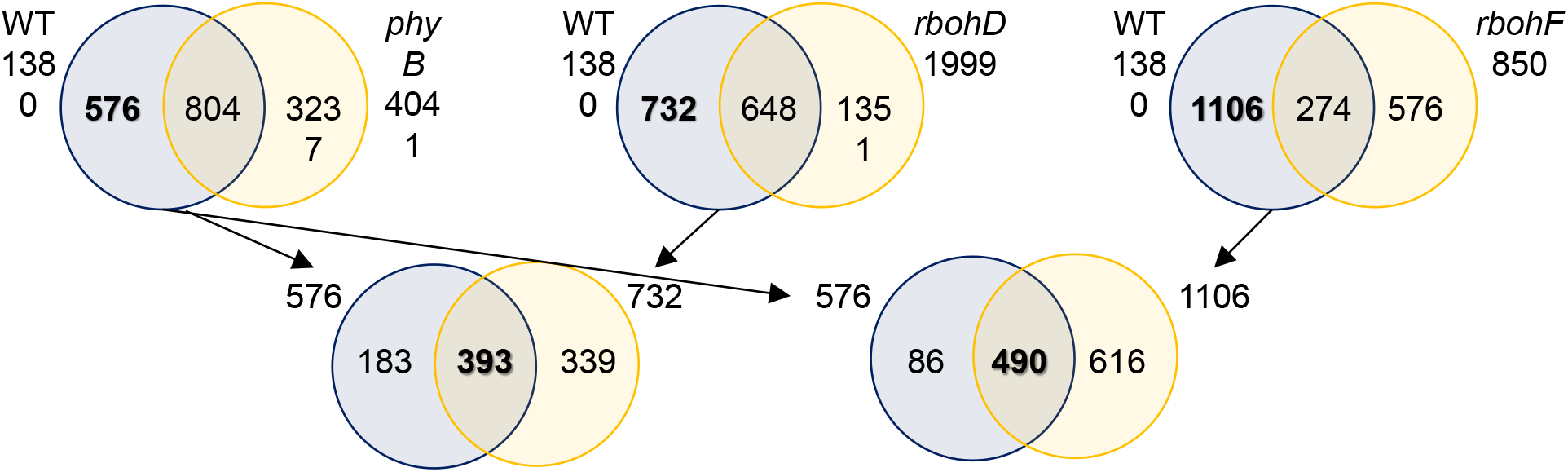
PhyB, RBOHD and RBOHF regulate the expression of thousands of transcripts in response to excess red light stress. Venn diagrams of the overlap between transcript altered in their expression in WT, *phyB*, *rbohD*, and *rbohF* plants in response to treatment with 10 min of excess red light. These Venn diagrams were used to deduce the 353 transcripts, shown in Fig. 2, induced by red light and are co-regulated by phyB, RBOHD and RBOHF. Abbreviations: phyB, phytochrome B; rbohD, respiratory burst oxidase homolog D; rbohF, respiratory burst oxidase homolog F; WT, wild-type.

**Figure S4.**
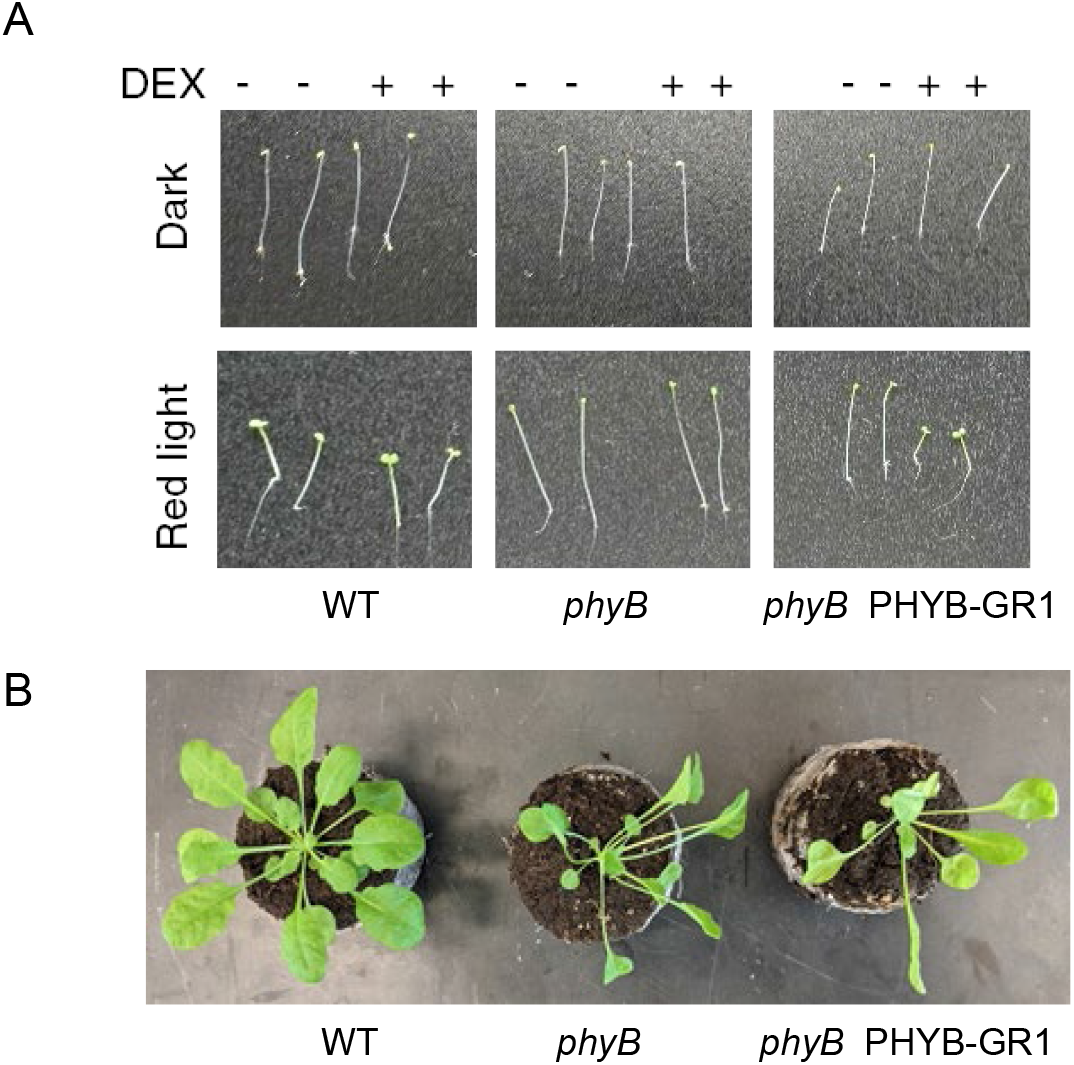
*phyB PHYB-GR1* phenotype in response to different illumination regimes and application of DEX. (A) Representative images of seedlings of WT, *phyB* and *phyB PHYB-GR1* seedlings grown on plates in dark or under constant red light without or with DEX. (B) Representative image of WT, *phyB* and *phyB PHYB-GR1* adult green plants, used for ROS measurement in Fig. 4A. Abbreviations: DEX, dexamethasone; GR1, Glucocorticoid Receptor; phyB, phytochrome B; ROS, reactive oxygen species; WT, wild-type.

**Figure S5.**
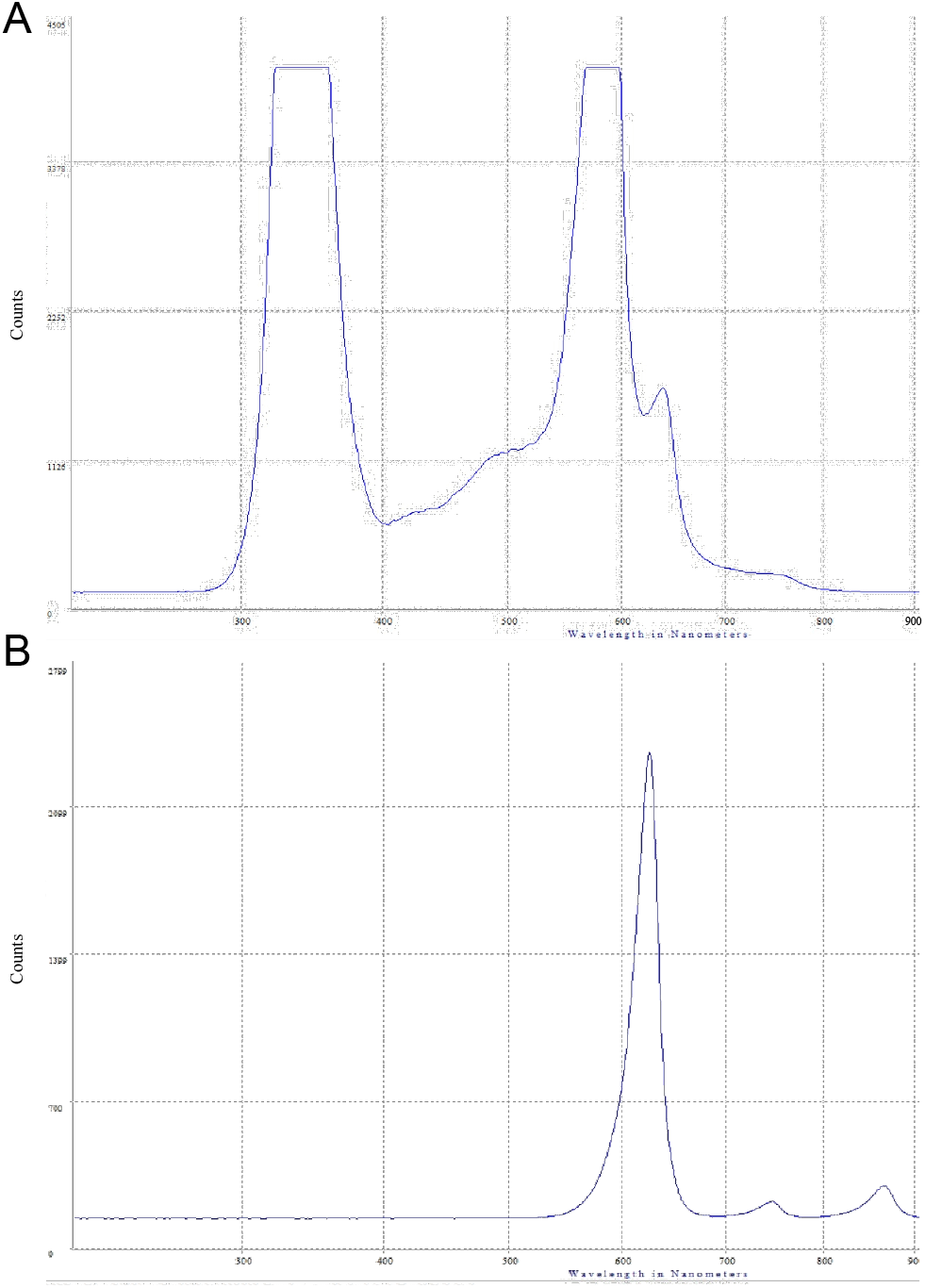
Wavelength spectrum of LED light sources used in this study. (A) Wavelength spectrum of white LED light source used for 10 min excess white light treatments. (B) Wavelength spectrum of red LED light source used for 10 min excess red light treatments. *Abbreviation:* LED, light emitting diode.

